# A systemic whole-plant change in redox levels accompanies the rapid systemic response of Arabidopsis to wounding

**DOI:** 10.1101/2020.11.04.366732

**Authors:** Yosef Fichman, Ron Mittler

## Abstract

Reactive oxygen species (ROS) play a key role in regulating plant responses to different abiotic stresses, wounding and pathogen attack. In addition to triggering responses at the tissues directly subjected to stress, ROS were recently shown to mediate a rapid whole-plant systemic signal, termed the “ROS wave”, required for inducing a state of systemic acquired acclimation, or systemic wound response. However, whether the ROS wave that spreads from the local tissues subjected to wounding to the rest of the plant triggers alterations in redox levels, is mostly unknown at present. Here, using a genetically-encoded reporter for cellular glutathione redox changes, roGFP1, we show that the wounding-induced systemic ROS wave in *Arabidopsis thaliana* is accompanied by a rapid systemic wave of cytosolic redox oxidation, termed a “redox wave”. The ROS wave may therefore trigger changes in redox levels in systemic leaves that in turn can trigger transcriptional, metabolic and proteomic changes resulting in acclimation and/or systemic wound responses.

**One sentence summary:** The wounding-induced reactive oxygen species (ROS) wave is accompanied by a systemic whole-plant redox response.

## Introduction

Reactive oxygen species (ROS) play a central role in the regulation of plant responses to different developmental signals, abiotic stresses, wounding and pathogen attack (*e.g*., Chang et al., 2004; Fryer et al., 2004; Mhamdi and Van Breusegem, 2018; Kollist et al., 2019). One of the major routes by which ROS cause an alteration in gene expression in response to many of these stimuli is a change in the redox state of different proteins (Mittler, 2017; Huang et al., 2019). Such proteins could include transcriptional regulators, phosphatases, kinases and different pumps and channels that link ROS signaling to a variety of different cellular responses and control plant acclimation and defense (Mittler, 2017; Huang et al., 2019). In addition to triggering responses at the tissues directly subjected to abiotic stress, ROS were recently shown to regulate rapid whole-plant systemic responses that occur at tissues not directly subjected to the stress, inducing a state of systemic acquired acclimation (SAA), or systemic wound response (SWR) (*e.g*., Miller et al., 2009; Szechyńska-Hebda et al., 2010; Suzuki et al., 2013; Gilroy et al., 2016; Fichman et al., 2019; Zandalinas et al., 2019; Zandalinas et al., 2020a). The ROS-dependent systemic signaling pathway mediating these processes, termed “the ROS wave” (Miller et al., 2009), was recently shown to be regulated by respiratory burst oxidase homolog proteins (RBOHD and RBOHF) and to be mediated through the vascular bundles of plants (Zandalinas et al., 2020b). However, whether the ROS wave that spreads from the local tissues subjected to stress to the rest of the plant triggers alterations in redox levels, is mostly unknown at present.

## Results and Discussion

The development of genetically-encoded reporters for cellular and glutathione redox changes, and their application in plants led to major advancements in the study of redox and ROS signaling in recent years (*e.g*., Schwarzländer et al., 2008; Meyer and Dick, 2010; Rosenwasser et al., 2010; Schwarzländer et al., 2016; Exposito-Rodriguez et al., 2017; Lim et al., 2019; Nietzel et al., 2019; García-Quirós et al., 2020; Haber and Rosenwasser, 2020). These reporters have however been primarily used in conjunction with confocal microscopy, limiting their application to the detection of redox changes in specific cells, tissues and organs. In contrast, whole-plant detection of redox changes in mature plants grown in soil has been limited. We recently developed a method for live whole-plant imaging of ROS levels in soil-grown plants and used it to study the ROS wave in wild type plants and different mutants responding to different stimuli (*e.g*., Devireddy et al., 2020; Fichman et al., 2020; Zandalinas et al., 2020a). Although changes in ROS levels (detected as accumulation of oxidized DCF in cells; Fichman et al., 2019) were demonstrated to accompany the systemic response of plants to different stimuli and these changes were shown to result in metabolic and transcriptomic changes that drove SAA or SWR (*e.g*., Suzuki et al., 2013; Zandalinas et al., 2019; Zandalinas et al., 2020a), it is not known whether a systemic whole-plant redox response also accompanies this rapid signaling process. To test this possibility, we studied cytosolic roGFP1-expressing plants (Jiang et al., 2006; Meyer et al., 2007; Schwarzländer et al., 2008; Supplementary material Methods) subjected to a local injury stimuli (Figures 1 and 2). The choice of roGFP1 was based on its highly reduced state at the cytosol which provides a low background in unstressed plants, and the choice of wounding was a result of not wanting to impact the roGFP1 sensor by treatments such as heat or high light stress that may alter its activity. As shown in Figure 1A, wounding of a single leaf of a wild type (Col-0) plant, grown in soil, resulted in the triggering of a systemic ROS wave response, imaged by the accumulation of oxidized DCF fumigated as H2DCFDA into whole-plants (Fichman et al., 2019). As shown in Figure 1B and Supplementary Movie 1, wounding of a single leaf of a cytosolic-expressing roGFP1 transgenic plant (Jiang et al., 2006), grown in soil, resulted in local and systemic changes in the redox state of the roGFP1 probe, evident by raw changes in Excitation/Emission at 420nm/520nm (for oxidized roGFP1) and Excitation/Emission of 480nm/520 nm (for reduced roGFP1) fluorescence. To quantify these changes and to calculate degree of oxidized roGFP1 in local and systemic tissues in response to wounding, we next determined the levels of 420nm/520nm (for oxidized roGFP1) and 480nm/520 nm (for reduced roGFP1) fluorescence in whole plants fumigated with 5 mM hydrogen peroxide (H_2_O_2_; to induce roGFP1 oxidation), or 5 mM Dithiothreitol (DTT; to induce roGFP1 reduction), for 15 min, and calculated the ratio of 420/480 nm for these treatments (Figure 2A). Using these ratios we then determined the 420/480 nm ratio and degree of roGFP1 oxidation in local and systemic leaves of the control and wounded plants shown in Figure 1 (Figure 2B and Supplementary Movies 2 and 3). The results shown in Figures 1B, 2 and Supplementary Movies 1-3, reveal that wounding of a single Arabidopsis leaf results in the oxidation of the cytosolic roGFP1 probe in both local and systemic leaves, and that these changes correspond to the changes recorded with the H2DCFDA probe (Figure 1A). Similar degrees of roGFP oxidation were recently reported for whole-plant imaging of chloroplastic roGFP2-expressing plants subjected to light stress (Haber and Rosenwasser, 2020), further supporting our findings and demonstrating that the degree of increase in roGFP oxidation in whole plants upon enhanced ROS accumulation is in the range of 5-10% (Figure 1B).

**Figure 1.**
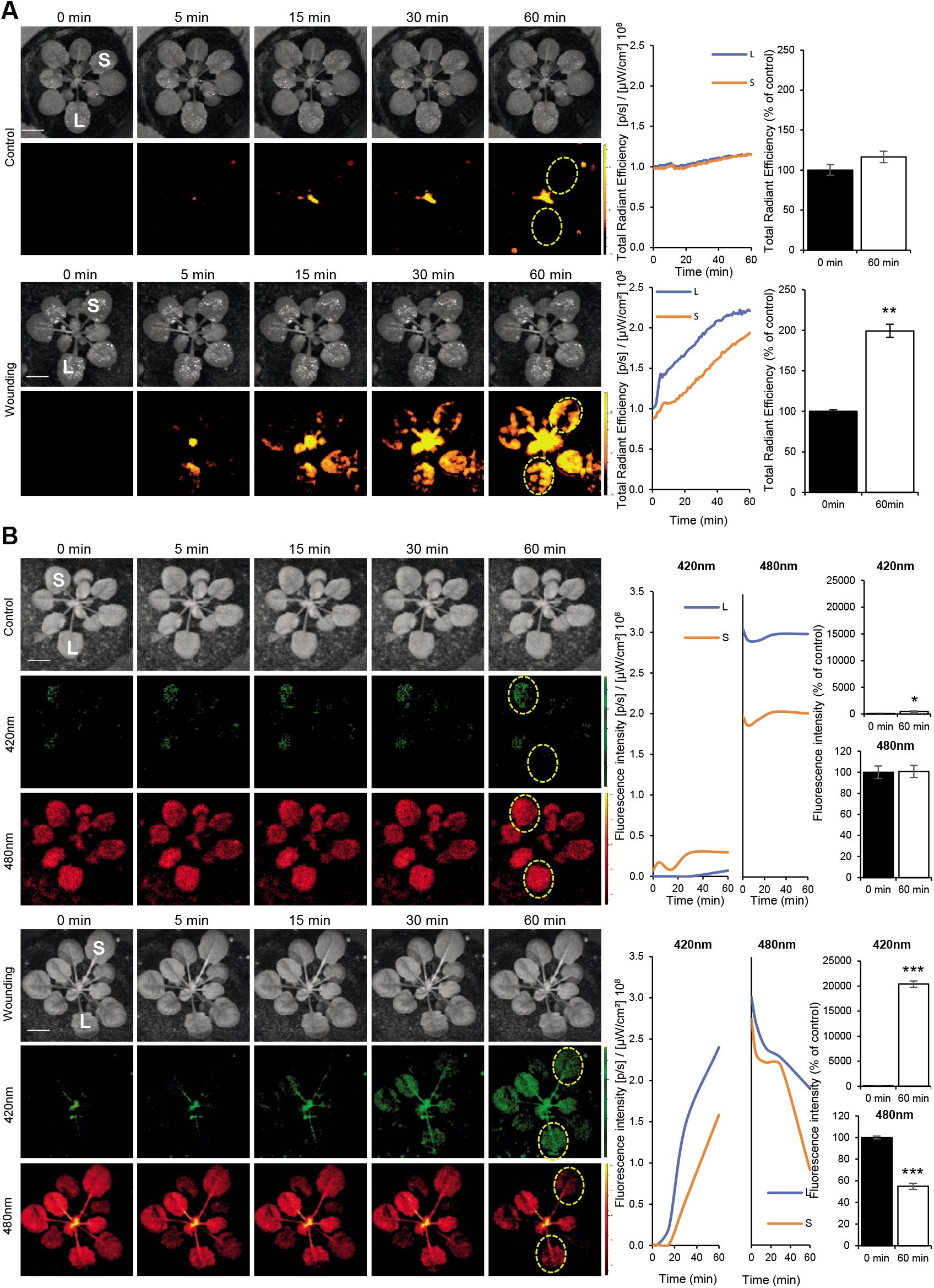
Imaging of ROS and redox levels in whole plants subjected to wounding. **A.** Representative time-lapse images of whole-plant ROS levels (indicated by DCF oxidation) in wild type *Arabidopsis thaliana* (Col-0) plants, untreated (control, top), or wounded (wounding, bottom; applied to leaf L only), are shown on left; representative line graphs showing continuous measurements of ROS levels in local (L) and systemic (S) leaves over the entire course of the experiment (0 to 60 min) are shown at the middle (ROIs used to generate the line graphs are indicated with dashed yellow ovals on left); and statistical analysis of ROS levels in local and systemic leaves at 0 and 60 min is shown on right. **B.** Same as in **A.**, but for transgenic plants overexpressing the roGFP1 protein at the cytosol (Jiang *et al*., 2006). Fluorescence was measured at excitation/emission (ex/em) of 420nm/520 nm for oxidized roGFP1 (middle, in green; oxidized) and at ex/em of 480nm/520 nm (bottom, in red; reduced). Experiments were repeated 3 times. Student t-test, SE, N=6, *P< 0.05, **P < 0.01, ***P < 0.005. Scale bar indicates 1 cm. *Abbreviations used:* DCF, 2’,7’-Dichlorofluorescein; ex/em, excitation/emission; L, local; S, systemic; roGFP1, reduction-oxidation sensitive green fluorescent protein 1; ROI, region of interest.

**Figure 2.**
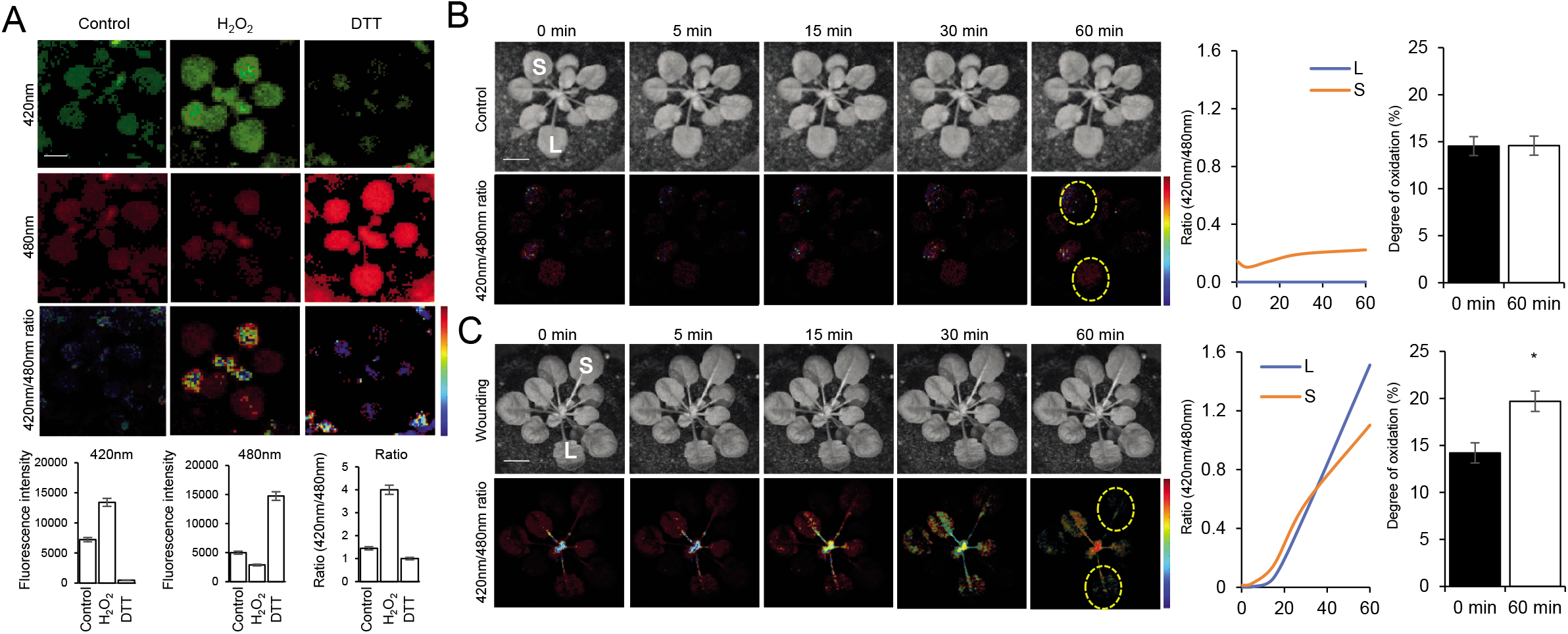
Whole-plant ratiometric fluorescence measurements of cytosolic roGFP1-expressing plants subjected to wounding. **A.** Representative images of whole-plant roGFP1 florescence (top; 420 and 480 nm excitations are in green and red, respectively), and statistical analysis of 420nm, 480nm and 420nm/480nm ratio (bottom) of untreated cytosolic roGFP1 plants (Control), or cytosolic roGFP1 plants subjected to a 15 min fumigation with 5 mM hydrogen peroxide (H_2_O_2_) or 5mM Dithiothreitol (DTT). Ratios of florescence intensities obtained from the H_2_O_2_ and DTT treatments were used for range normalization of the degree of roGFP1 oxidation (similar to Schwarzländer et al., 2008) shown in B. **B.** Representative time-lapse images (left) and line graphs of 420/480 nm ratio for the entire time course (ROIs used to generate the graphs are shown in the images as dotted yellow ovals; middle), and bar graphs showing degree of roGFP1 oxidation for the 0 and 60 min time points calculated using the normalization range obtained in A (right), generated for the control untreated plants shown in Figure 1B. **C.** Same as in **B.**, but for the wounded plants shown in Figure 1B. Experiments were repeated 3 times. Student t-test, SE, N=6, *P < 0.05. Scale bar indicates 1 cm. *Abbreviations used:* DTT, Dithiothreitol; L, local; S, systemic; roGFP1, reduction-oxidation sensitive green fluorescent protein 1; ROI, region of interest.

Our findings that wounding of a single leaf is accompanied by a systemic wave of ROS production (Fichman et al., 2019; Figure 1A), as well as a change in cytosolic redox levels (a “redox wave”; Figures 1B, 2; Supplementary Movies 1-3) could therefore provide an initial clue to how the systemic ROS wave response alters the levels of metabolites and transcripts in systemic tissues causing an enhanced state of SAA or SWR (Miller et al., 2009; Szechyńska-Hebda et al., 2010; Suzuki et al., 2013; Gilroy et al., 2016; Fichman et al., 2019; Zandalinas et al., 2019; Zandalinas et al., 2020a, 2020b). Interestingly, the roGFP1 probe remained oxidized for at least 60 min following wounding, showing that the ROS wave response is turned “on” for at least 1 hour following wounding (Figure 1B). This finding is in agreement with our recent findings that the systemic ROS wave response to excess light stress remains “on” for at least 3 hours following the initial application of the 10 min high light stress treatment to the local leaf (Devireddy et al., 2020). Further studies are of course needed to identify the different regulatory proteins altered by the ROS and redox waves in response to a local stimuli and to tie their altered activity to the systemic response they cause. One potential regulator, recently identified during systemic responses to light stress, is MYB30 (Fichman et al., 2020). Because changes in MYB30 protein oxidation were shown to alter its DNA binding activity, MYB30 could be a potential regulator that links changes in redox levels in systemic tissues to transcript expression and SAA (Tavares et al., 2014; Fichman et al., 2020). In addition to demonstrating that the ROS wave is accompanied by a change in redox levels in local and systemic tissues (Figures 1 and 2; Supplementary Movies 1-3), our study also demonstrates that at least some genetically-encoded reporters can be imaged in whole-plants grown in soil, using a sufficiently sensitive apparatus (Fichman et al., 2019; Figures 1, 2, and Supplementary Movies 1-3). This finding opens the way for further studies of ROS and redox signaling in whole plants grown in soil, and perhaps even to large-scale phenotyping studies.

## Acknowledgments

The roGFP1 seeds were obtained from the Arabidopsis Biological Resource Center (ABRC). This work was supported by funding from the National Science Foundation (MCB-1936590, IOS-1932639, and IOS-1353886), and the University of Missouri. We apologize to all authors of papers not mentioned in this manuscript due to space limitations.

