## Supplementary material Methods for "A systemic whole-plant change in redox levels accompanies the rapid systemic response of Arabidopsis to wounding"

### Methods (in support of Figure 1 and 2)

#### Plant Material and Growth conditions

*Arabidopsis thaliana* Columbia-0 plants and roGFP1 expressing plants (c-roGFP1-7, CS25418, ABRC; Jiang *et al.,* 2006) were grown in peat pellets (Jiffy International, Kristiansand, Norway) under controlled conditions of 10hr/14hr light/dark regime, 50 µmol $m^{-2}$ $s^{-1}$, 21°C for 4 weeks. Wounding was performed as described in Fichman *et al.,* 2019 by puncturing a single leaf, with 18 dressmaker pins (Singer, Murfreesboro, TN, USA).

#### ROS imaging with $H_2$DCFDA and roGFP1 fluorescence measurements

Imaging was performed using the IVIS Lumina S5 platform (PerkinElmer, Waltham, MA, USA). ROS imaging with 2',7'-Dichlorofluorescein (DCF; Excitation/Emission 480nm/520nm; Millipore-Sigma, St. Louis, MO, USA) was performed as described in Fichman *et al.* , 2019, 2020, Fichman and Mittler 2020, Zandalinas *et al.,* 2020a, 2020b; Devireddy *et al.,* 2020. Briefly, plants were fumigated for 30 min with 50 µM $H_2$DCFDA solution (50 mM phosphate buffer, pH 7.4, 0.01% v/v Silwet L-77), wounded on a single leaf and placed in the imager. Images of DCF fluorescence were acquired every 60 s for 60 min. For roGFP1 fluorescence, plants were wounded and placed in the imager. Oxidized roGFP1 was detected by Excitation/Emission of 420nm/520nm and reduced roGFP1 by Excitation/Emission of 480nm/520 nm. Images were acquired every 60 s for 60 min. Images were analyzed using Living Image 4.7.2 software (PerkinElmer) as described in Fichman *et al.* , 2019, 2020, Zandalinas *et al.,* 2020a, 2020b; Devireddy *et al.,* 2020. roGFP1 oxidized/reduced ratio images were calculated on a pixel-by-pixel basis, using MATLAB RatioImage application (Advanced Imaging Center; https://www.mathworks.com/matlabcentral/fileexchange/50340-ratioimage). Data was normalized by obtaining oxidized and reduced ranges of roGFP1, as follows: For oxidized, plants were fumigated with 5 mM $H_2O_2$ for 15 min and then imaged, and for reduced roGFP1, plants were fumigated with 5 mM Dithiothreitol (DTT) for 15min and then imaged (Figure 2A). Ratio values were normalized between 4 to 1 (oxidized roGFP1 to reduced roGFP1), based on intensity values of $H_2O_2$ and DTT fumigation, similar to Jiang *et al*., 2006 and Schwarzländer *et al*., 2008. Normalized ratios were used for calculating degree of oxidation (Schwarzländer *et al*., 2008), with the following equation:

$$OxD\mathrm{roGFP1} = \frac{R-Rred}{\frac{I480ox}{I480red}(Rox-R)+(R-Rred)}$$

In which, OxDroGFP1 is degree of oxidation for roGFP1, R is fluorescence ratio of 420/480 nm, Rred is the ratio of 5mM DTT reduced form, Rox is the ratio of 5 mM $H_2O_2$ oxidized form and I480ox and I480red are the intensities at 480 nm for the 5 mM H2O2 oxidized and 5 mM DTT reduced forms (Schwarzländer *et al*., 2008).

**Statistical Analysis**

Data obtained from the different analyses and calculations described above, and from the different images using the Living Image software, was statistically analyzed using Microsoft Excel. Statistical significance was determined by two-tailed student t-test. At least 3 plants were used for the experiments and results are presented as mean ± SE (*P< 0.05, **P < 0.01, ***P < 0.005).

**Supplementary movie legends.**

**Supplementary Movie 1.** Time-lapse video imaging of raw whole-plant changes in cytosolic roGFP1 fluorescence (Oxidized roGFP1 was detected by Excitation/Emission of 420nm/520nm and reduced roGFP1 by Excitation/Emission of 480nm/520 nm) in response to wounding applied to the local leaf only (indicated with a yellow circle). Plants were wounded as described in Fichman *et al.,* 2019. Supporting Figure 1.

**Supplementary Movie 2.** Time-lapse video imaging of local and systemic changes in ratio of oxidized to reduced (Excitation/Emission of 420nm/520nm and Excitation/Emission of 480nm/520 nm) cytosolic roGFP1 in control plant (no treatment). Supporting Figure 2.

**Supplementary Movie 3.** Time-lapse video imaging of local and systemic changes in ratio of oxidized to reduced (Excitation/Emission of 420nm/520nm and Excitation/Emission of 480nm/520 nm) cytosolic roGFP1 in a wounded plant. Supporting Figure 2.
